# The Epithelial-to-Mesenchymal Transition Supports HCMV Infection and Biosynthesis in Mammary Epithelial Cells Through Two Distinct Mechanisms

**DOI:** 10.1101/2024.08.12.607657

**Authors:** Preethi Golconda, Mariana Andrade-Medina, Rachel Matrenec, Alan McLachlan, Adam Oberstein

## Abstract

Human cytomegalovirus (HCMV) infects a wide range of cell types in the body, including a variety of epithelial cell types. Despite the significance of epithelial cells during infection, HCMV has been difficult to study in epithelial cells. In this study, we examined HCMV infection in mammary and prostate epithelial cell lines, finding that the virus establishes a semi-permissive, biosynthetically abortive state. Building on previous work, we hypothesized that shifting epithelial cells to a mesenchymal cell state would restore HCMV biosynthesis and progeny production. To test this hypothesis, we induced epithelial-to-mesenchymal transition (EMT) using TGF-β and the EMT-transcription factor (EMT-TF) SNAIL. We found that shifting strongly epithelial cell lines to a mesenchymal cell state shifted HCMV infection from a semi-permissive to fully permissive state. This effect appeared to involve two distinct mechanisms: EMT-sensitive enhancement of viral entry and EMT-sensitive enhancement of viral mRNA translation. Although the precise mechanisms remain elusive, our findings identify the epithelial-mesenchymal cell state axis as an important regulator of HCMV infection and provide new insights into how cellular differentiation states influence viral replication. They also raise the possibility that the EMT pathway, a fundamental pathway involved in development and cancer metastasis, could regulate HCMV infection *in-vivo*, potentially contributing to viral persistence or pathogenesis in epithelial tissues.

## Introduction

Amongst the eight known human herpesviruses, human cytomegalovirus (HCMV) is remarkable in that it has an unusually broad cellular tropism ^1,2^. HCMV is able to infect a variety of somatic cells in the body, including fibroblasts, epithelial cells, endothelial cells, and smooth-muscle cells; as well as myeloid cells and neuronal cells ^3,4^. Consequently, many organ systems are affected during HCMV disease, including liver, kidney, lung, intestine, oral mucosa, skeletal muscle, and vascular endothelium ^1,5^ . In the absence of immunological control, HCMV can cause a variety of pathologies ranging from mild to lethal. These include cytomegalic inclusion disease, inflammation of the gut, liver, or lung; hearing loss in infants, and graft rejection in transplant patients ^5–7^.

The virus is also suspected to contribute to a range of chronic pathologies, including immune senescence, atherosclerosis, and cancer ^8–15,15–19^. Although the mechanisms underlying these phenomena are poorly understood, the long time-frame required for the development of these pathologies suggests they could be collateral consequences of viral persistence. HCMV causes a life-long persistent infection in humans, with cycles of reactivation and latency recurring intermittently over decades, allowing ample opportunity for the virus to acutely or permanently dysregulate the biology of host cells.

Accumulating evidence suggests that HCMV can persist locally at particular physiologic sites in the body, in addition to the myeloid compartment, the primary reservoir thought to harbor latent viral genomes. For example, HCMV is persistently shed from healthy, seropositive, individuals via bodily fluids such as urine ^5,20^, saliva ^20–24^, and breast milk ^25–27,27,28^. Additionally, viral antigens and nucleic acids have been found in tumor tissue of the mammary gland ^10^, prostate gland ^29,30^, ovary ^31^, and brain ^9^; leading some to suspect that the virus directly contributes to tumorigenesis ^11,15,19,32–35^.

Epithelial cells are important players in both the normal tropism of HCMV, serving portals into and out of the body ^1^ , and HCMV pathogenesis, which often presents in epithelial tissues ^4,36,37^ . However, despite the important role epithelial cells play during HCMV infection, relatively little is known about how HCMV interacts with epithelial cells. Few cell culture models exist for examining HCMV in non-transformed cells and each epithelial cell type, despite having common cell biological features, could interact with HCMV in a different manner, due cell type specific differences in biochemical function and/or epigenetics. Furthermore, both laboratory preparations of HCMV easily lose epithelial tropism when expanded in commonly utilized fibroblast cell lines ^38–42^, which has presented a technical barrier to studying HCMV in epithelial cells.

Lastly, we recently found that one of the most well studied epithelial models for studying HCMV *in-vitro*, the ARPE-19 retinal pigment epithelial cell line, when cultured under typical conditions, assumes a mesenchymal cell state and behaves like a fibroblast cell line, rather than an epithelial cell line, in experimental infection assays ^43^. We also discovered that ARPE-19 cells lack a canonical type-I interferon response to HCMV ^44^. Both of these observations suggest that ARPE-19 cells, at least in the subconfluent, undifferentiated state, may not provide physiologically relevant insights into the behavior of HCMV in epithelial cells.

These findings led us to seek alternative cell culture models for studying HCMV in epithelial cells, including the cell lines RWPE-1 and MCF10A. RWPE-1 cells were derived from normal prostate tissue after immortalization with the E6 and E7 proteins from human papillomavirus 18 ^45^, while MCF10A cells were derived from mammary tissue and are spontaneously immortalized ^46^. Both cell lines are non-transformed (i.e. they will not form tumors in animal models) and both retain strong epithelial characteristics and normal cell-type specific biochemistry ^45,46^. We have found that both cell lines are susceptible to HCMV infection ^43^. However, to our surprise, experimental infection of either line with HCMV results in a semi-permissive infection, where viral biosynthesis is aborted and cell free infectious progeny production is restricted ^43^. During this work, we also noted that productive HCMV infection appeared to correlate mesenchymal cell state features and negatively correlated with epithelial cell state features, leading us to suspect that HCMV biosynthesis might be sensitive to cellular pathways feeding into the epithelial-mesenchymal cell-state axis.

Thus, we hypothesized that HCMV biosynthesis might be restored in semi-permissively infected epithelial cells after trans-differentiation to a mesenchymal cell state. To test this hypothesis, we have assessed the impact of initiating an epithelial-to-mesenchymal transition (EMT) in prostate and mammary epithelial cells on HCMV biosynthesis and infectious progeny production. We find that the HCMV infectious cycle can be shifted from a semi-permissive to fully-permissive state by driving epithelial cells to a mesenchymal cell state using either chemical or genetic stimuli. The mechanism behind this phenomenon appears to involve cell state sensitive translation of essential viral mRNAs, whose lack of production in some epithelial cells can limit viral biosynthesis. Furthermore, this restriction can be overcome by driving epithelial cells to a mesenchymal cell state by stimulating the cellular EMT pathway. Lastly, we find that viral entry is enhanced in mammary epithelial cells after mesenchymal trans-differentiation, suggesting that susceptibility to infection may be dynamically regulated in some epithelial cell types.

Our work raises the possibility that the EMT pathway may be involved in controlling HCMV biosynthesis *in-vivo* and we hypothesize that under particular conditions HCMV may establish a biosynthetically poised state in epithelial cells that can be restored to a productive state by epithelial trans-differentiation and/or other overlapping developmental pathways. Such a mechanism could contribute to HCMV pathogenesis or persistence in epithelial tissues.

## Material and Methods

### Cell Lines, Culture Conditions, and Viruses

MRC-5 human embryonic lung fibroblasts and ARPE-19 adult retinal pigment epithelial cells were obtained from the American Type Culture Collection and grown in DMEM supplemented with 10% fetal bovine serum, 1 mM sodium pyruvate, 2 mM Glutamax (Gibco), 10 mM Hepes pH 7.4, 0.1 mM MEM Non-Essential Amino Acids (Gibco), 100 units/mL Penicillin G, and 100 μg/mL Streptomycin Sulfate. MCF10A cells were cultured in DMEM supplemented with 5% horse serum, 10 mM Hepes pH 7.4, 20 ng/ml EGF, 10 ug/ml Insulin, 1 nM Forskolin, 500 ng/ml Hydrocortisone, 100 units/ml Penicillin G, and 100 μg/ml Streptomycin Sulfate. RWPE-1 cells were cultured in K-SFM (Gibco) supplemented with 25 units/ml Penicillin G and 25 μg/ml Streptomycin Sulfate. HEK293FT and HEK293-GP2 cells were from ThermoFisher and Dr. Garry Nolan (Stanford University), respectively. HCMV strain TB40/E- BAC4 ^47^ was reconstituted by electroporation into ARPE-19 cells, expanded on 15 cm dishes of ARPE-19 cells, and concentrated by ultracentrifugation over a 20% sorbitol cushion. Virion-containing pellets were resuspended in PBS containing 7% sucrose and 1% BSA, aliquoted, and stored at -80° C until use. Virus preparations were titered on both MRC-5 and ARPE-19 cells using an HCMV infectious unit assay, as previously described^43^.

### Vector and Stable Cell Line Construction

Stable cell lines expressing doxycycline inducible GFP or Snail were created using pLVX-TetOne-Puro (Takara Bio Inc., Shiga, Japan). A lentiviral transfer vector expressing LgBit was prepared by ligating a synthetic gene fragment (Twist Bioscience, San Francisco, USA) into retroviral vector pLXSN (Takara Bio Inc., Shiga, Japan). A lentiviral transfer vector expressing a translational fusion between HCMV pp65 and HiBit was prepared by first PCR amplifying pp65 from TB40/E-BAC4 BAC DNA using primers containing restriction site overhangs, and then ligating the digested pp65 DNA fragment and a double-stranded, annealed oligo encoding the HiBit-peptide, into pLV-EF1a-IRES-Puro (a gift from Tobias Meyer; Addgene plasmid #85132; RRID: Addgene_85132). A stable, complementing, cell line for loading pp65:HiBit into HCMV virions was then created by transducting ARPE-19 cells with pLV-EF1a-pp65:HiBit-IRES-Puro lentiviruses and selecting using puromycin (see below). ARPE-19 and MCF10A cells doubly transduced with pLV-EF1a-LgBit-IRES-Puro and pLVX-TetOne-Snail-Neo, and selected with G418 and Puromycin (both from Sigma-Aldrich, St. Louis, MO, USA) were used as target cells for Nanoluciferase entry assays (see below). All plasmids were validated by sanger or nanopore sequencing prior to use.

Lentiviral vectors were produced by co-transfecting each lentiviral transfer vector with the packaging plasmids pCMV-dR8.91 and pCMV-VSV-G into HEK293FT cells. Plasmids were mixed in a 12:12:1 (vector:dR8.91:VSV-G) ratio by mass and mixed with an empirically optimized amount of branched polyethyleneimine (Sigma-Aldrich, St. Louis, MO, USA) before adding to cells. Lentivirus-containing supernatants were collected at 48 and 72 h post-transfection and concentrated over a 20% sorbitol cushion using ultracentrifugation. Lentivirus pellets were resuspended in PBS containing 7% Sucrose and 1% BSA. Retroviral vectors were produced in a similar manner by transfecting retroviral transfer vectors into HEK293-GP2 cells.

Transductions were performed by empirically determining the volume of lentivirus allowing 60 to 70% of the cells to survive puromycin selection; an effective MOI of approximately 1.0. 5 μg/mL hexadimethrine bromide (“polybrene”; Sigma-Aldrich, St. Louis, MO, USA) was added during transduction. Forty-eight hours after transduction, selective media containing 2 μg/mL puromycin was added until non-transduced cells were completely killed, at which point transduced cells were maintained in 1 μg/mL puromycin for expansion. G418 prepared in 25 mM Hepes-NaOH pH 7.4 was used at 2 mg/ml for selection.

To construct plasmids for preparation of Southern probes, the intronless HCMV genes UL79 and UL95 were PCR-amplified from high-molecular weight bacmid DNA (strain TB40/E-BAC4) and ligated into pLVX-Tetone-PURO containing a modified multiple cloning site.

### Quantitative reverse transcription polymerase chain reaction (*q*RT-PCR)

RNA was isolated using Trizol reagent (Ambion) followed by purification using Qiagen Rneasy columns. Contaminating DNA was removed using Turbo Dnase (Thermo Fisher Scientific) and 1 ug total RNA was reverse-transcribed in a 20 ul reaction volume using a Superscript VILO cDNA synthesis kit (Invitrogen). *q*RT-PCR reactions were performed in 384-well plates in a 10 ul reaction volume by mixing 0.7 - 1.7 ul of 10-fold diluted cDNA from each sample with 0 – 1.0 ul H20, 5 ul 2x Power SYBR Green Master Mix (Applied biosystems) and 3.3 ul 1.2 uM PCR primers ([primer]final = 0.4 uM). qRT-PCR reactions were performed using Power SYBR Green Master Mix (Applied biosystems) and data were collected on a Viia7 digital PCR machine (Life Technologies). Data were analyzed using the ΔΔCT method^48^ with cyclophilin A (PPIA) as the reference gene. Primers were designed using either QuantPrime^49^ or GETPRime^50^. *q*RT-PCR primer sequences are listed in Supplementary Table S1.

### RNA sample preparation

For RNA-seq, ARPE-19 and MCF10A cells were infected at a multiplicity of 3 or 5 infectious units/cell, respectively. Cells from three independent infections were collected in Trizol reagent (Applied Biosystems) at 6, 24, 48, 72, 96, and 120 hpi and total RNA was isolated using an RNeasy RNA isolation kit (Qiagen). DNA was removed using Turbo DNase (Thermo Fisher Scientific) and RNA was quantified and quality-controlled using an Agilent TapeStation 4200. Sequencing libraries were prepared by the UIC College of Medicine Genomics Research Core Facility using a CORALL Total RNA-seq Kit with Ribosomal RNA depletion (Lexogen) and sequenced on an Illumina NovaSeq DNA sequencer in paired-end, rapid mode (50 bp x 2).

### RNA-seq data analysis

Concatenated human/HCMV fasta and annotation (.gtf) files were created for mapping and feature counting by combining sequences and annotations from gencode human genome Release 24 (GRCh38.p5) and TB40E-BAC4 (EF999921.1). HCMV long non-coding RNA’s were annotated in TB40/E by querying the TB40/E genome with the corresponding sequences from HCMV strain Merlin (NC_006273.2) with tblastn ^51^. RNA-sequencing fastq files were pseudo-aligned to the homo sapiens transcriptome and converted to pseudo-counts using Kallisto version 0.45.0 ^52^. Gene-level read counts were summed from the Kallisto transcript pseudocounts using tximport ^53^. The count matrices for all runs were joined and normalized for transcript length (within sample normalization) and compositional bias (cross-sample normalization) using the GeTMM ^54^ procedure and the R-package limma ^55,56^. Data were plotted using the R-package ^57^ ggplot2 ^58^.

### Western blotting and immunofluorescence staining of E-cadherin

For western analysis cell monolayers were collected in RIPA buffer (50 mM Hepes, 150 mM NaCl, 1 % NP-40, 0.5 % sodium deoxycholate, 0.1 % SDS, 5 ug / ml aprotinin, 10 ug / ml leupeptin, 1 mM PMSF, pH 7.4), sonicated, and cleared by centrifugation. Protein concentration was determined using the BCA assay (Pierce) and equal amounts of total protein were separated by SDS-PAGE. Proteins were transferred to PVDF membranes and blocked with 5 % BSA in HBST (50 mM Hepes pH 7.4, 150 mM NaCl, 0.05 % tween-20). Primary antibodies were diluted in 1 % BSA and rocked at 4 °C overnight. Primary antibodies were detected using HRP-conjugated secondary antibodies and ECL Prime (Cytiva). Antibodies and dilutions are listed in Supplementary Table S2.

For immunofluorescence staining, monolayers were grown in collagen-treated, glass bottom, 96-well tissue culture plates (Greiner), infected with HCMV as described above, and allowed to progress to the indicated times post infection. Monolayers were then fixed with 2% paraformaldehyde/PBS for 15 min at RT, quenched with 50 mM NH_4_Cl_2_ in PBS for 30 min, permeabilized with 0.5 % triton-X in PBS for 10 min at RT, washed three times with PBS, and then blocked with 5% human serum (Sigma; H3667) for 1 h at room temperature. Monolayers were then stained with a panel of anti-HCMV monoclonal antibodies (a generous gift from Dr. Thomas Shenk, Princeton University) diluted in 5% human serum and monolayers were rocked at 4 °C overnight. Primary antibodies were detected using Alexa Fluor-conjugated secondary antibodies (ThermoFisher).

The directly conjugated anti-IE1 antibody was created by labeling 100 ug MAB8131 (Millipore-Sigma) with CF568 using a commercial amine-reactive, antibody labeling kit (Sigma; MX568S100-1KT); using the manufacturers instructions.

Antibodies and dilutions are listed in Supplementary Table S2.

### Experimental Infections

For experiments using doxycycline induction, MCF10A cell lines (i.e. MCF10A-iGFP or MCF10A-iSnail) were treated with 1 ug / ml doxycycline (Sigma) in 15 cm dishes for two days, split one-to-three after 48 of treatment, then treated for two additional days. Cells were then counted and seeded into 24-well plates at 2 x 10^5^ cells per well (-dox), or 2.5 x 10^5^ cells per well (+dox), and maintained for 24 additional hours at 37°C and 5 % CO_2_. Cells were infected at the indicated multiplicities by adding an appropriate HCMV inoculum to the cell culture supernatant for 2 h and returning the dishes to a standard tissue culture incubator set at 37°C and 5 % CO_2_. After the infection period, cells were washed one time with warm PBS and returned to the tissue culture incubator in complete medium. MOIs were calculated using titers obtained from ARPE-19 cells.

### HCMV Entry Assays

With the exception of the nanoluciferase entry experiments, an IE1-immunofluorescence assay was used as a surrogate for viral entry. The assay was performed as previously described ^43^. Cells were seeded into 24-well plates in biological triplicate for each MOI and infected with HCMV as described above (see Methods; “Experimental Infections”). At 24 hpi, monolayers were fixed in cold MeOH and stained for IE1 antigen using monoclonal antibody 1B12 ^59^. Cells were counterstained with DAPI to visualize nuclei and the percentage of infected cells in each condition was calculated using fluorescence imaging by dividing the number of IE1-positive nuclei by the total number of DAPI-positive nuclei using CellProfiler ^60^ v2.1.1 software.

Nanoluciferase internalization assays were performed by first generating a stock of HCMV with pp65:HiBit loaded into its tegument, by infecting 15 cm plates of pLV-EF1a-pp65:HiBit-IRES-Puro-ARPE-19 cells (see “Plasmids” above) with HCMV TB40/E-BAC4. The resulting virus preparation was titered on ARPE-19 cells and used to infect target ARPE-19 or MCF10A-iSnail cells stably transduced with pLXSN-LgBit.

To perform the assay MCF10A-iSnail-LgBit cells were pretreated with doxycycline or vehicle for 4 days, then reseeded at equal density into 96-well tissue culture plates. ARPE-19-LgBit cells were used as a control, as were cells pretreated for one hour with 10 ug/ml heparin (Sigma, USA) or 3% Cytogam (CSL Behring, USA). On the day of the assay, target cells were first incubated on ice for 20 min to halt virus internalization. TB40/E-pp65:HiBit virus was then added to the cell supernatant at a multiplicity of 1 IU/cell and allowed to attach to cells for one hour on ice. Unbound virus was then removed, and monolayers were washed once with ice-cold PBS before shifting the temperature to 37 °C by placing them in a tissue culture incubator. After a 15 min internalization period, cells were lysed in Nano-Glo Live Cell Reagent (Promega), and luminescence was detected with Synergy2 Microplate Reader (BioTek). Relative internalization was determined by first subtracting the background signal from uninfected control wells for each treatment and cell type, averaging the background subtracted luminescence signal of three biological replicates, and normalizing to the cell type with maximal average luminescence signal.

### HCMV Infectious Progeny Assays

Both infectious unit (IU) assays and plaque assays were used to assess HCMV infectious progeny. The infectious unit assay was performed as previously described ^43^. At the indicated time post-infection, cell supernatants were collected, cleared of detached cells by centrifuging at 300 × g for 5 min at RT, aliquoted, and frozen at −80 °C until all time points were collected. Once all time points were harvested, supernatants were thawed (1 × freeze–thaw cycle), serially diluted in DMEM/10% FBS in 96-well plates, and adsorbed onto confluent monolayers of MRC-5 fibroblasts. Twenty hours later, reporter fibroblasts were fixed with cold MeOH, stained for HCMV immediate early-1 (1E1) antigen using monoclonal antibody 1B12 ^59^, imaged using a Keyence BZ-X710 Inverted Microscope, and titers were calculated as follows. Wells with 20% or less infected nuclei were selected to avoid saturation effects and IE1-positive nuclei in each image were counted using CellProfiler v2.1.1 ^60^l. The mean titer and standard deviation for each condition was calculated by multiplying the number of IE1-positive nuclei on each image by (1/dilution factor), (1.0 mL/infection volume) in milliliters, and a well factor. The well factor was the total well surface area (acquired from the plate manufacturer’s engineering specification) divided by the image area (calculated using a calibration slide and the imageJ (Version 1.52) ^61^ function “set scale”). Four images/well (technical replicates) were sampled from three independent infections (biological replicates). Infectious units per mL (IU/mL) were determined by averaging the three technical replicates for each biological replicate (using the equation described above), converting each average to a titer, and then calculating the average and standard deviation of the biological replicates.

Plaque assays were also used in some experiments, to confirm infectious progeny were competent to complete all phases of infection. For this, cell supernatants were harvested, stored, and serially diluted identically to the IU assay described above. Diluted supernatants were then adsorbed onto confluent monolayers of MRC-5 fibroblasts in 24-well plates in a tissue culture incubator. After two hours, supernatants were aspirated, monolayers were washed 1x with PBS, and a 1:1 mixture of 1.0 % low-melting point agarose and 2x DMEM/20% FBS maintained at 42 °C was added to each well. Agarose was left to solidify at RT in a tissue culture hood for 1 h and then returned to a tissue culture incubator. Plaques were scored manually using an inverted phase contrast microscope at 10 and 14 days post-infection. Plaque-forming units per milliLiter (PFU/mL) were calculated by taking the average and standard deviation from three independent wells for each condition and multiplying by 1/dilution factor and 1.0 mL/infection volume.

### Southern Blotting

MCF10A-iSnail cells were cultured in the presence or absence of 1ug/ml doxycycline for 4 days. One day prior to infection, MCF10A-iSnail (-dox) and MCF10A-iSnail (+dox), and ARPE-19 cells were seeded into 10 cm dishes at 2x10^6^, 3.5x10^6^, or 2x10^6^ cells/well, respectively. The next day, cells were infected with HCMV at 3.0 MOI (MCF10A-iSnail -dox) or 0.4 MOI (MCF10A-iSnail +dox; ARPE-19) to equalize the percentage of infected cells in each cell type. The viral inoculum was incubated with cells for two hours in a tissue culture incubator. After the infection period, monolayers were washed one time with PBS and returned to the tissue culture incubator in complete medium for the indicated times. Infected cells treated with the viral polymerase inhibitor phosphonoformic acid (PFA) were used as a negative control. At 24, 48, 72, and 120 hpi total DNA was isolated from cells using a Proteinase K/Phenol extraction procedure ^62^. At each time point, cells were washed with cold phosphate buffered saline (PBS), scraped into PBS, centrifuged at 4°C for 10 min at 1500 x*g*, and resuspended in lysis buffer (10 mM Tris-HCL pH 8.0, 0.1 M EDTA, 0.5 % (w/v) SDS, 20 ug/ml RNase A) at 5x10^6^ cells/per mL. Lysates were then treated with 100 ug/mL Proteinase K (Sigma) for 3 hours at 50 °C followed by phenol extraction and ethanol precipitation. Southern blotting was performed by digesting 10 ug total DNA from each sample with BamHI (NEB, Ipswich, MA) overnight at 37 °C. DNA was then concentrated by ethanol precipitation, separated on a 1 % agarose gel, transferred to nitrocellulose by upward capillary transfer ^63^, and probed with P-32 labeled dsDNA probes prepared from plasmids encoding the HCMV UL79 and UL95 genes. Blots were detected using a phosphorous screen and Typhoon Phosphorimager (Cytiva Life Sciences).

### Measurement of Cell Associated Viral Genomes by quantitative PCR (*q*PCR)

For DNA *q*PCR analysis, cells were collected in Trizol Reagent (Sigma). Total DNA was extracted from the aqueous phase by ethanol precipitation followed by two washes in 0.1 M Sodium Citrate in 10 % ethanol (pH 8.5), one wash with 75% ethanol, and pelleting by centrifugation. DNA was resuspended in 8 mM Hepes-NaOH and RNA was removed by treatment with 10 ug/ml RNAse A for 1 h at 37°C, followed by phenol/chloroform extraction, and a second round of ethanol precipitation. The final DNA pellet was resuspended in 8 mM NaOH followed by the addition of 1/10 volume of 1M HEPES-NaOH. *q*PCR reactions were performed using Power SYBR Green Master Mix (Applied biosystems) and data was collected on a Viia7 digital PCR machine (Life Technologies). Viral genomes were detected using primers to HCMV *UL44* and the number of HCMV genomes per cell were calculated as 2^-ΔCt of UL44 compared to the cellular reference gene cyclophilin A (PPIA). Samples treated with 100 uM Ganciclovir (Sigma) served as negative controls for viral DNA replication.

## Data Availability

Kallisto pseudo counts, study metadata, count matrices, and code used to perform normalization, and data visualization are available at the following public github repository: XXX. RNA-sequencing data will be deposited in the NCBI Gene Expression Omnibus (GEO). DNA plasmids and cell lines are available from the corresponding author upon reasonable request.

## Results

### EMT induction in epithelial cells supports HCMV replication and infectious progeny production

Given the correlation we previously observed between productive infection and mesenchymal cell-state features ^43^, we hypothesized that initiating an EMT in epithelial cells might convert them from a semi-permissive, biosynthetically stalled state, to a fully permissive, productive, biosynthetic state. To test this hypothesis, we trans-differentiated MCF10A and RWPE-1 cells from an epithelial to mesenchymal cell-state using chemical or genetic stimuli, challenged them with an epithelial-tropic preparation of HCMV strain TB40/E-BAC4 ^41–43,47^, and monitored the effect on HCMV biosynthesis and infectious progeny production. As a first test, we chose chemical treatment with the EMT-inducing cytokine transforming growth factor beta (TGF-β), a well-characterized fibrogenic cytokine widely employed to induce mesenchymal characteristics in a variety of epithelial cell types ^64–68^. MCF10A and RWPE-1 cultures were treated with 1 ng/ml TGF-β or vehicle for seven days, replated at equal density, infected with HCMV at a multiplicity of 5 infectious units per cell (IU/cell), and assayed for cell-free infectious progeny production at 120 hours post infection (hpi) (Fig. 1). MCF10A cells transitioned from a cobblestone, epithelial, morphology to a flattened, spindle-shaped, morphology (Fig. 1A); were infected at high efficiency (Fig. 1B); and produced 35-fold more infectious progeny than untreated or vehicle treated cells (Fig. 1C). RWPE-1 cells showed little change in morphology, but produced 12-fold higher levels of infectious progeny after treatment with TGF-β (Fig. 1C), suggesting that EMT-induction enhances HCMV biosynthesis in both cell lines.

**Figure 1.**
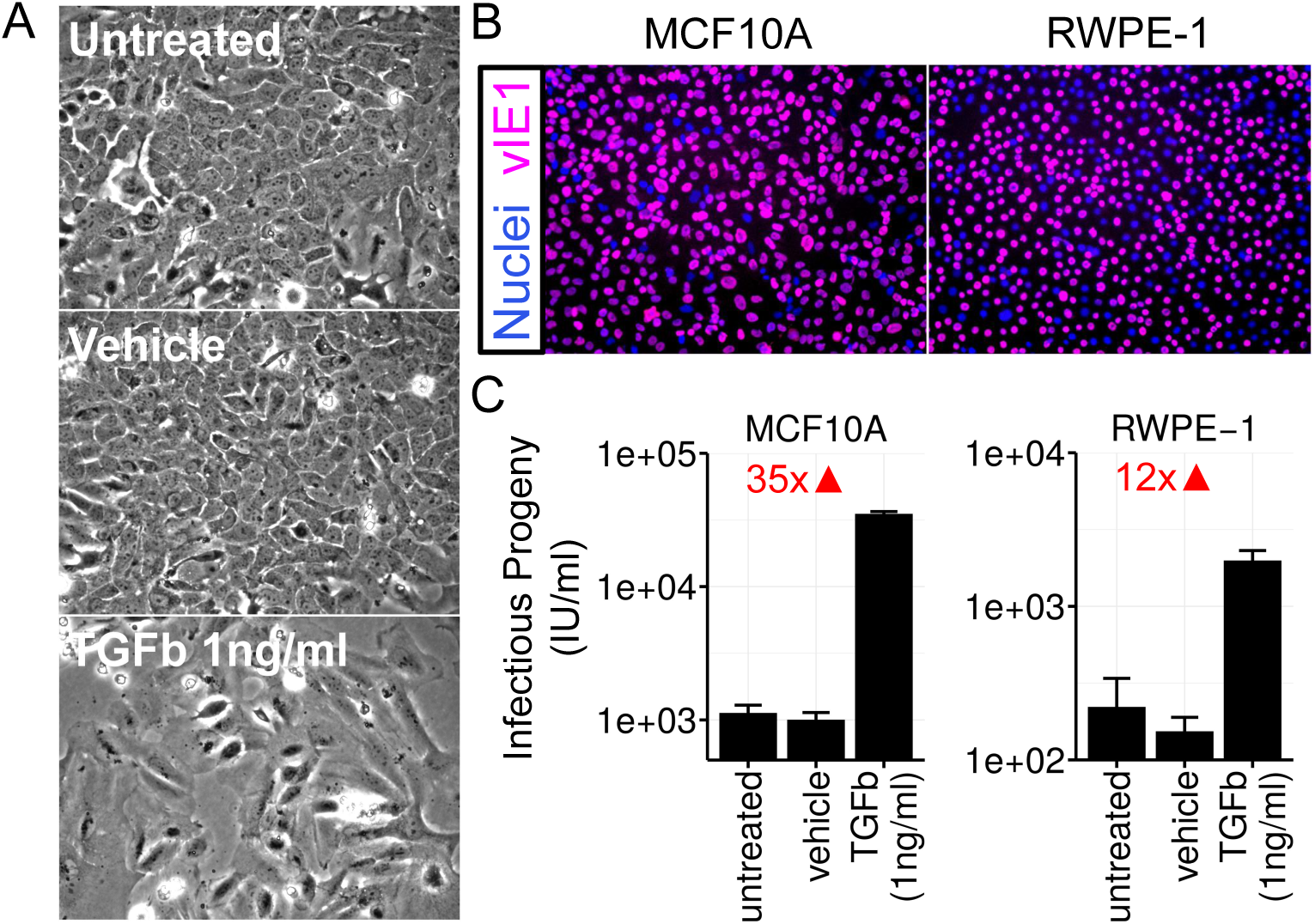
Induction of EMT in MCF10A and RWPE-1 Cells using TGF-β Enhances HCMV Infectious Progeny Production. (A) Phase-contrast images depicting morphological changes induced in MCF10A cells treated with 1 ng/ml TGF-β, vehicle (PBS), or left untreated for 7 days. Cells were split at day 4 and allowed to reach confluency prior to imaging. (B) Assessment of viral entry using indirect immunofluorescence staining of the HCMV immediate early-1 (IE1) protein at 16 hpi. Infections were performed at 5 MOI and monolayers were prepared for microscopy (see “methods”). Magenta = IE1; Blue = dapi/nuclei. (C) HCMV Infectious progeny production at 120 hpi, in the presence or absence of TGF-β treatment. Cells were pre-treated with TGF-β or control for 7 days, re-plated at equal density, and infected with an epithelial-tropic stock of TB40/E-BAC4 at 5 MOI. Infectious progeny production at 120 hpi was assessed using an infectious unit assay (see “methods”) on MRC5 fibroblasts.

TGF-β has broad, pleiotropic, effects that can be cell type specific. Therefore, we also assessed the impact of genetic induction of EMT on HCMV biosynthesis. The EMT-inducing transcription factors (EMT-TFs) SNAIL, SLUG, ZEB1, ZEB2, TWIST1, and TWIST2 control EMT during development, wound healing, and cancer cell metastasis ^69,69–74^. Ectopic expression of individual EMT-TFs is sufficient to initiate an epithelial-to-mesenchymal transition in a variety of epithelial cell types and the EMT-TF SNAIL is an important downstream effector of TGF-β in epithelial cells ^68,75–77^. However, long-term expression of EMT-TFs in epithelial cells can affect viability and or cell-division ^78^. Therefore, we constructed a genetic system to conditionally express SNAIL, or the control, green-fluorescent protein (GFP), in MCF10A mammary epithelial (Fig. 2A) cells. MCF10A cell lines cells were transduced with “all-in-one” inducible lentiviruses expressing SNAIL or GFP, stable transductants were selected using puromycin, and the resulting populations (referred to as “MCF10A-iSNAIL” or “MCF10A-iGFP”) were challenged with HCMV before or after induction of EMT. Fully permissive ARPE-19 cells transduced with the same GFP-expressing lentivirus (ARPE-19-iGFP) served as an additional control.

**Figure 2.**
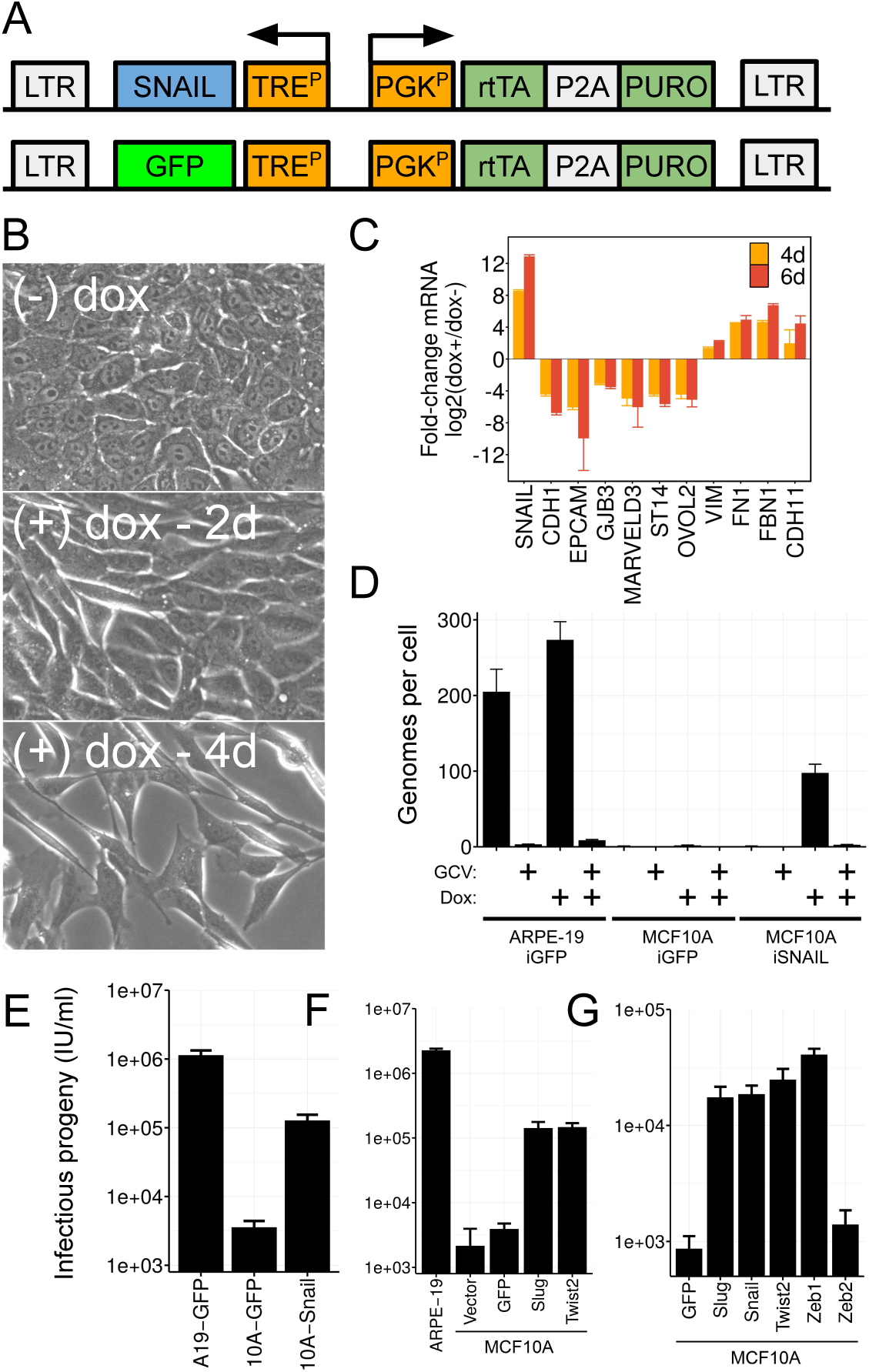
Induction of EMT in MCF10A and RWPE-1 Cells using EMT-transcription factors (EMT-TFs) Enhances HCMV Replication and Infectious Progeny Production. (A) Diagram of inducible lentivirus vectors constructed to transitenly express the EMT-TF SNAIL, or a GPF control protein in MCF10A cells. SNAIL or GFP were inserted downstream of the a tetracycline inducible promoter in the vector backbone pLVX-Tetone-PURO, and stable cells lines were created by lentiviral transduction and selection with puromycin. (B) Phase-contrast images depicting morphological changes induced in MCF10A cells treated for 2 and 4 days of 1 ug/ml doxycycline. MCF10A cells transition from a cobblestone, epithelial morphology, to a spindly, mesenchymal morphology indicative of EMT. (C) Relative mRNA levels of epithelial (*CDH1, EpCAM, GJB3, MARVELD3, ST14, OVOL2)* and mesenchymal (*VIM, FN1, FBN1, CDH11*) marker genes, as well as *SNAIL*, measured by *q*RT-PCR. MCF10A-iSnail cells were treated with vehicle (water) or 1 ug/ml doxycycline for 4 or 6 days prior to RNA-extraction. Data represent the mean +/- SD from three biological replicates assayed in technical triplicate. (D) HCMV viral DNA production across control (ARPE19-iGFP, MCF10A-iGFP) and EMT-induced cells (MCF10A-iSnail). Total DNA was isolated from cell populations infected with TB40/E-BAC4 at 5 MOI (MCF10A) or 3 MOI (ARPE-19) at 120 hpi and the number of HCMV genomes per cell were measured using quantitative PCR to the HCMV *UL44* gene. Untreated control populations served to control for the effects of dox on HCMV genome replication, and populations treated with ganciclovir confirmed the specificity of the PCR signal. (E) HCMV infectious progeny production from Snail/EMT-induced and control populations (same as in panel D). All populations were pretreated with 1 ug/ml dox, re-plated at equal densities, and infected at 5 MOI with HCMV TB40/E-BAC4. Progeny were collected at 120 hpi and titered using an IE1-infectious unit assay on MRC5 fibroblasts. (F and G) Same as panel E, except that constitutively expressing lentiviral vectors were used instead of dox-inducible vectors. Data in all infectious progeny assays represent the average +/- SD of three independent biological replicates per condition.

In the absence of the inducer doxycycline (dox), stable MCF10A-iSNAIL derivatives exhibit a parental-like, cobblestone morphology (Fig. 2B). Treatment with doxycycline, potently induced SNAIL mRNA expression (Figs. 2C and S2) and progressively induced a mesenchymal phenotype in MCF10A cells, as assessed by cobblestone-to-spindle morphological transition (Fig. 2B) and gene expression changes consistent with EMT (i.e. inhibition of epithelial marker genes and induction of mesenchymal marker genes; Figs. 2C and S1). We have also previously shown that E-cadherin and EpCAM steady-state protein levels are potently reduced in dox-treated MCF10A-iSNAIL cells, while Vimentin proteins levels are increased ^43^; verifying the EMT-inducing effects of our genetic system.

To assess the effects of EMT on HCMV infection, we treated MCF10A-iSNAIL cells for four days in the presence or absence of 1 ug/ml doxycycline, re-plated each culture at an equal cell density (Fig. S2), infected them with HCMV at an equal multiplicity or infection. We then monitored viral DNA accumulation (Fig. 2D) and infectious progeny production (Fig. 2E). HCMV-infected MCF10A-iGFP and ARPE19-iGFP cells controlled for unanticipated metabolic effects of doxycycline, which can affect some assays ^79–81^. Infected ARPE19-iGFP cells produced approximately 200 - 250 HCMV ganciclovir-sensitive genomes per cell and 1x10^6^ cell-free infectious units per milliliter (IU/ml) at 120 hpi, indicative of robust HCMV replication. Parental-like MCF10A-iGFP cells maintained a small number of cell-associated viral genomes (∼ five/cell) and produced little-to-no infectious progeny (∼ 5x10^3^ IU/ml). This level of infectious progeny results from residual input inoculum adsorbed to MCF10A cells, which we have observed previously in single-step growth assays ^43^. Induction of SNAIL in MCF10A cells partially rescued both viral DNA replication and infectious progeny production, with the number of cell-associated viral genomes per cell increasing 20-fold and infectious progeny production increasing approximately 50-fold compared to MCF10A-iGFP cells (Figs. 2D and 2E).

To determine if the observed effects on HCMV were due specifically to SNAIL or a general effect of initiating the EMT program, we assessed whether additional EMT-TFs could similarly enhance infectious progeny production in MCF10A cells. For this, we created an isogenic set of stable MCF10A sublines constitutively expressing one of the six EMT-TFs or GFP, and measured infectious progeny production after infection with HCMV. Two independent experiments were performed, with independently transduced cell populations (Figs. 2F and 2G). SNAIL, SLUG, ZEB1, and TWIST2 supported infectious progeny production, while ZEB2 did not. We could not establish a stable, constitutive Twist1-expressing MCF10A cell line, and therefore it remains unclear whether TWIST1 phenocopies other EMT-TFs with respect to HCMV infection. Since multiple EMT-TFs were found to support HCMV progeny production, the effect of EMT-TFs on the HCMV infectious cycle appears to be an effect of the EMT program in general, rather a specific effect of one particular EMT-TF.

Overall, these data show that the HCMV infectious cycle is sensitive to the epithelial-mesenchymal cell state of epithelial cells. They also support the notion that one or more features of the epithelial cell-state program may be intrinsically restrictive to HCMV infection and that induction of a mesenchymal state via the EMT can relieve such restrictions.

### MCF10A cells contain two nested restrictions to HCMV infection that are overcome by EMT induction

To gain insight into the mechanism(s) of HCMV restriction in MCF10A cells, we first asked whether HCMV gene expression was initiated normally in MCF10A cells. In permissive cells, HCMV gene expression follows a well-characterized temporal cascade, progressing from immediate-early (IE), to early (E), to late (L) gene expression. This cascade serves as a master control script for virion biosynthesis, producing mRNAs encoding a variety of viral transcription factors, enzymes, and structural proteins. To monitor the HCMV gene expression cascade in MCF10A cells, total RNA was isolated from HCMV-infected MCF10A cells at multiple time points after infection and subject to RNA-sequencing analysis. Absolute viral RNA levels were assessed by placing transcript counts on a common, normalized, reads-per-kilobase-per-million (RPKM) scale (see Methods). RNA-sequencing measurements from infected ARPE-19 cells were used as permissive controls. Transcription from the HCMV genome showed remarkable similarity in both kinetics and expression level, between ARPE-19 and MCF10A cells (Fig. 3A, 3B, and S3). However, viral RNA levels were suppressed between 2 and 10-fold in MCF10A cells at late time points (> 24 hpi) (Fig. 3B, S3, and S4). Nonetheless, all kinetics classes of transcripts accumulated to near-productive (ARPE-19) levels in MCF10A cells, with many late viral transcripts achieving 500 - 1000 RPKM (9 - 10 log2 RPKM), an absolute expression level similar to highly expressed cellular mRNAs (e.g. compare *UL84*, *RNA5.0*, *UL94*, or *UL99* with *PPIA* in Fig. S4). These data suggested that transcription from the HCMV genome occurrs fairly normally in non-productively infected MCF10A cells.

**Figure 3.**
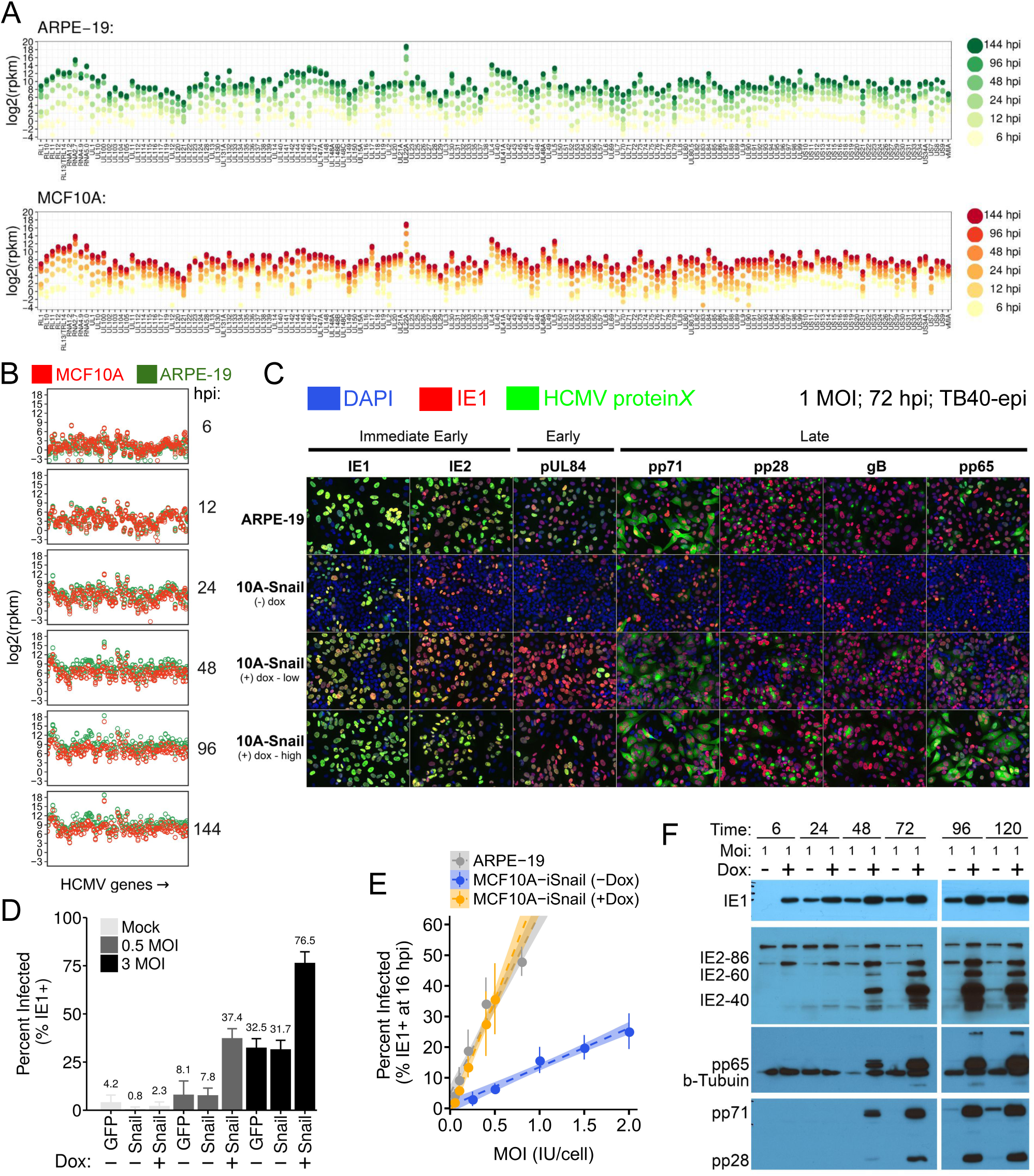
MCF10A cells contain two nested restrictions to HCMV infection that are overcome by EMT induction. (A) Relative mRNA levels of canonical HCMV transcripts in HCMV-infected ARPE-19 and MCF10A cells. Absolute mRNA levels were quantified using RNA-sequencing at different times after infection. Data points represent the average mRNA levels in log2 reads-per-kilobase-per-million (RPKM) of three independent infections, at each time point. Cells were infected at 3 MOI (ARPE-19) or 5 MOI (MCF10A) with TB40/E-BAC4 grown in ARPE-19 cells. (B) Direct comparison of HCMV transcript levels across infected ARPE-19 and MCF10A cells, binned by time point. Same data from panel A. (C) Immunofluorescence analysis of select HCMV proteins across productive ARPE-19 cells and MCF10A-iSnail cells in the presence and absence of dox treatment. All cells were infected at 1 MOI with HCMV TB40/E-BAC4 grown in ARPE-19 cells (“TB40-epi”) and prepared for immunofluorescence analysis at 72 hpi. Green signals are from indirect immunofluorescence staining against the viral proteins IE1. IE2, UL84, pp71, gB, and pp65. The red signal is from a second IE1 antibody directly conjugated to CF568 fluorescent dye and marks infected cells in each population. DAPI was used to counterstain cellular nuclei (blue). +dox - low = 0.1 ug/ml; +dox - high = 1 ug/ml. (D) HCMV IE1-entry assays used to assess relative entry of HCMV in MCF10A-iSnail cells in the presence or absence of EMT-induction, at low (0.5) and high (3.0) MOI. MCF10A-iGFP cells we used as a controls. (E) Relationship between the percentage of infected cells (assessed by IE1-assay) and multiplicity of infection (MOI). Trend lines represent a linear regression of the data from each cell type. (F) Western time course of HCMV proteins in MCF10A-iSnail cells, in the presence or absence of doxycycline. Both populations were infected at 1.0 MOI and lysates were harvested for immunoblot analysis at the indicated times.

Given that HCMV transcription was largely intact in MCF10A cells, we next assessed whether translation of viral proteins occurred normally in MCF10A cells. For this, we monitored HCMV protein expression using two methods: immunofluorescence analysis and immunoblotting. First, we monitored the expression of a panel of temporally expressed HCMV proteins (IE1 and IE2 = *immediate-early*; pUL84 = *early*; pp71, pp28, gB, and pp65 = *late*) in ARPE-19 and MCF10A-iSnail cells, in the presence or absence of low (100 ng/ml) or high (1 ug/ml) doses of doxycycline. To differentiate cells that failed to be infected, from those that were infected but abortive, samples were co-stained with a directly conjugated antibody to IE1 (Fig 3C and S5; red staining), in addition to indirectly conjugated antibodies directed to each temporal CMV antigen (Fig 3C and S5; green staining). Each cell-type was infected at a multiplicity of one infectious unit per cell (IU/cell) with an epithelial-tropic preparation of TB40/E-BAC4 (see “methods”). Productive ARPE-19 cells showed strong expression of all classes of viral proteins at 72 hpi (Fig. 3C) and 120 hpi (Fig. S5). In the absence of Snail/EMT induction, MCF10A cells showed a decreased frequency of IE1 expressing cells, as well as severely diminished numbers of IE1-positive (red) cells co-expressing early and late viral proteins (Figs. 3C and S5). Induction of EMT by expressing Snail increased the frequency of IE1-positive cells and restored early and late viral protein expression, phenocopying productively infected ARPE-19 cells (Figs. 3C and S5). The tegument protein pp65 appeared to be trapped in the nucleus in MCF10A cells (Figs. 3C and S5), suggesting a possible defect in pp65 trafficking in epithelial-state MCF10A cells.

By infecting MCF10A cells at different multiplicities, in the presence and absence of Snail/EMT, we confirmed that EMT increased the frequency of IE1-expressing cells by 4.7-fold and 2.3-fold, at low and high multiplicity, respectively (Fig. 3D and Fig. S6). Modeling of the relationship between the percentage of infected cells and the infection multiplicity showed that Snail/EMT induction converted MCF10A epithelial cells from a moderately susceptible state to a highly susceptible state similar to ARPE-19 cells (Fig. 3E). Immunoblot analysis of HCMV-infected MCF10A-iSnail cells confirmed the rescue of late protein expression in the presence of Snail/EMT and the restriction of early and late proteins in the absence of EMT (Fig. 3F). Similar results were observed in MCF10A cells expressing Snail from a constitutive promoter (Fig. S7).

These data suggest that EMT supports HCMV infection via two distinct mechanisms: one upstream of IE1 expression, possibly involving viral entry, and one downstream of IE1 expression, possibly involving EMT-sensitive translation of viral mRNAs.

### EMT Potentiates HCMV Attachment and Internalization in Mammary Epithelial Cells

Given the large effect of EMT on the frequency of IE1-expressing MCF10A cells, we suspected that HCMV entry was being modulated by the EMT. Therefore, we comparatively assessed HCMV attachment and internalization across MCF10A, MCF10A-EMT, and highly susceptible, control, ARPE-19 cells. PCR measurements of internalized HCMV genomes showed a high degree of variability across experiments. Therefore, we developed a luciferase-based assay to assess virus internalization (Fig. 4A). In this assay, the tegument protein pp65 is first loaded into HCMV virions by infecting ARPE-19 “producer” cells ectopically expressing a translational fusion between pp65 and a 1.3-kDa “HiBit” peptide from split nanoLuciferase ^82,83^. pp65:HiBit virions are then titered and used to infect target cells ectopically expressing the second half of nanoLuciferase, the 18-kDa LgBit protein ^82,83^, in the cytoplasm. Upon internalization, LgBit reassembles with pp65:HiBit, reconstituting nanoLuc luminescence, which is detected using a luminometer. Using this system, we assessed differential internalization of HCMV at 15 min post infection across ARPE-19, MCF10A-iSnail (-dox), and MCF10A-iSnail (+dox) cells. In the absence of doxycycline, MCF10A-iSnail cells consistently showed 2-fold lower luciferase levels than fully susceptible control ARPE-19 cells or MCF10A-iSnail cells treated with doxycycline to induce Snail/EMT (Fig. 4B). The luciferase signal in these assays was dependent on HCMV entry, as luminescence was reduced to background in the presence of anti-HCMV immunoglobulin (cytogam), which blocks viral entry.

**Figure 4.**
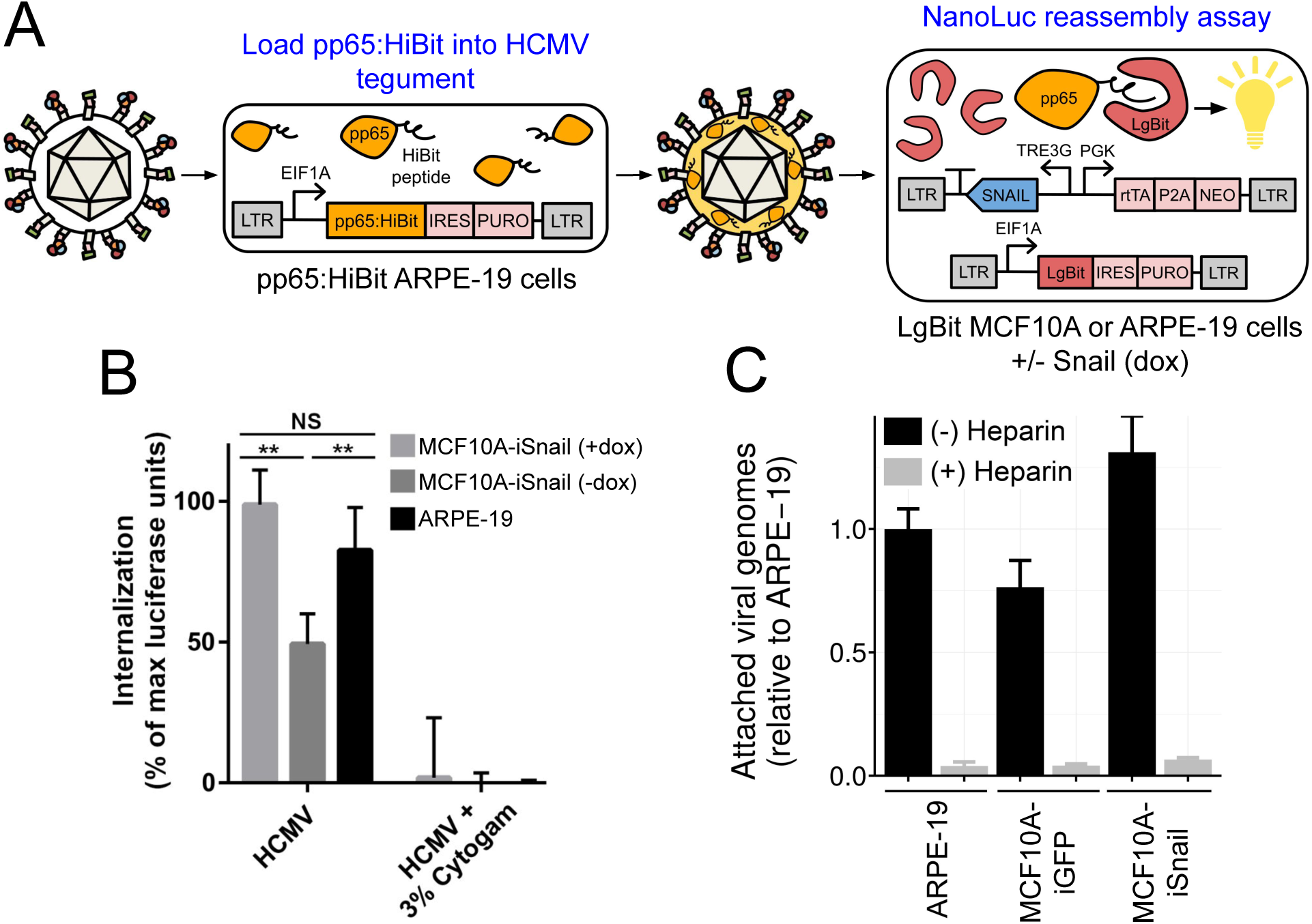
EMT Potentiates HCMV Attachment and Internalization in Mammary Epithelial Cells. (A) Principle of split-nanoLuciferase HCMV internalization assay. A fusion between the tegument protein pp65 and HiBit is loaded into HCMV virions by infecting complementing ARPE-19 “producer” cells. pp65:HiBit virions are then titered and used to infect target cells ectopically expressing LgBit in the cytoplasm. Upon internalization, LgBit reassembles with pp65:HiBit, reconstituting nanoLuciferase, whose enzymatic activity converts a substrate into a luminescent signal. (B) NanoLuc HCMV entry assays across highly susceptible ARPE-19 cells (positive control) and MCF10A-iSnail cells treated, or not treated, with doxycycline to induce EMT. Pre-treatment with anti-HCMV immunoglobulin (Cytogam) was used to assess signal specificity. Data represent the mean +/- SD of three independent replicates after background subtraction. (C) Assessment of HCMV attachment on ice to ARPE-19, MCF10A-iGFP, and MCF10A-iSnail cells treated with 1 ug/ml doxycycline. An equal volume of viral inoculum (equivalent to 1 MOI) was adsorbed onto each cell population on ice for 1 h, prior to extensive washing, and total DNA isolation. The relative amount of attached viral genomes were then measured by quantitative PCR. Data represent mean +/- SD of three independent biological replicates per condition. Soluble heparin sulfate, an attachment inhibitor, was used to control for signal specificity.

We also assessed possible effects of EMT on virus attachment by measuring the relative quantities of protease-resistant viral genomes adsorbed onto MCF10A-iGFP and MCF10A-iSnail cells treated with doxycycline. Compared to control ARPE-19 cells, MCF10A-iGFP cells adsorbed ∼ 30% fewer heparin-sensitive viral genomes, while MCF10A-iSnail cells adsorbed 30% more heparin-sensitive viral genomes (Fig. 4C).

These data suggest that MCF10A cells have a reduced capacity to both bind and internalize HCMV particles and that EMT potentiates viral entry by enhancing both of these processes.

### HCMV Biosynthesis and Infectious Progeny Production After Adjusting for Differential Viral Entry

The reduced susceptibility of epithelial-state MCF10A cell (i.e. MCF10A-iSnail -dox) could bias assays dependent on population averaging (e.g. bulk *q*RT-PCR or western analysis). Therefore, we performed an independent series of assays aimed at correcting the bulk-cell measurements, adjusting for the difference in viral entry observed across MCF10A cells before and after inducing EMT. To do so, we selected two sets of MOIs resulting in approximately equal frequencies of IE1-expressing cells at 24 hpi across ARPE-19, MCF10A-iSnail-dox, and MCF10A-iSnail +dox populations. These MOIs were: 0.4 / 3.0 / 0.4 (∼ 35 % infected) and 0.2 / 1.0 / 0.2 (∼ 20 % infected) for ARPE-19 / MCF10A-iSnail -dox / MCF10A-iSnail +dox cells, respectively. Using these “normalized” infection conditions, we re-assessed HCMV biosynthesis and infectious progeny production in MCF10A cells in the presence or absence of Snail/EMT induction (Figure 5). Similar to previous experiments using *equal MOIs*, viral DNA (Fig. 5A) and mRNA (Fig. 5B) production was slightly reduced in untreated/epithelial MCF10A-iSnail cells compared to dox-treated/mesenchymal MCF10A-iSnail cells. MCF10A-iSnail cells phenocopied fully productive ARPE-19 cells (Figs. 5A and 5B). However, steady-state viral protein levels were strongly suppressed in both indirect immunofluorescence (Figs. 5C and S8) and immunoblotting assays (Fig. 5D), suggesting that their production was sensitive to the epithelial-mesenchymal cell-state of the MCF10A cells. Lastly, infectious progeny production was restricted in MCF10A-iSnail cells in the absence of dox and restored to near-ARPE-19 levels in the presence of dox (Fig. 5E); again, suggesting that the ability to produce cell-free infectious virions could be modulated by the epithelial-mesenchymal cell state of the host cells exposed to HCMV.

**Figure 5.**
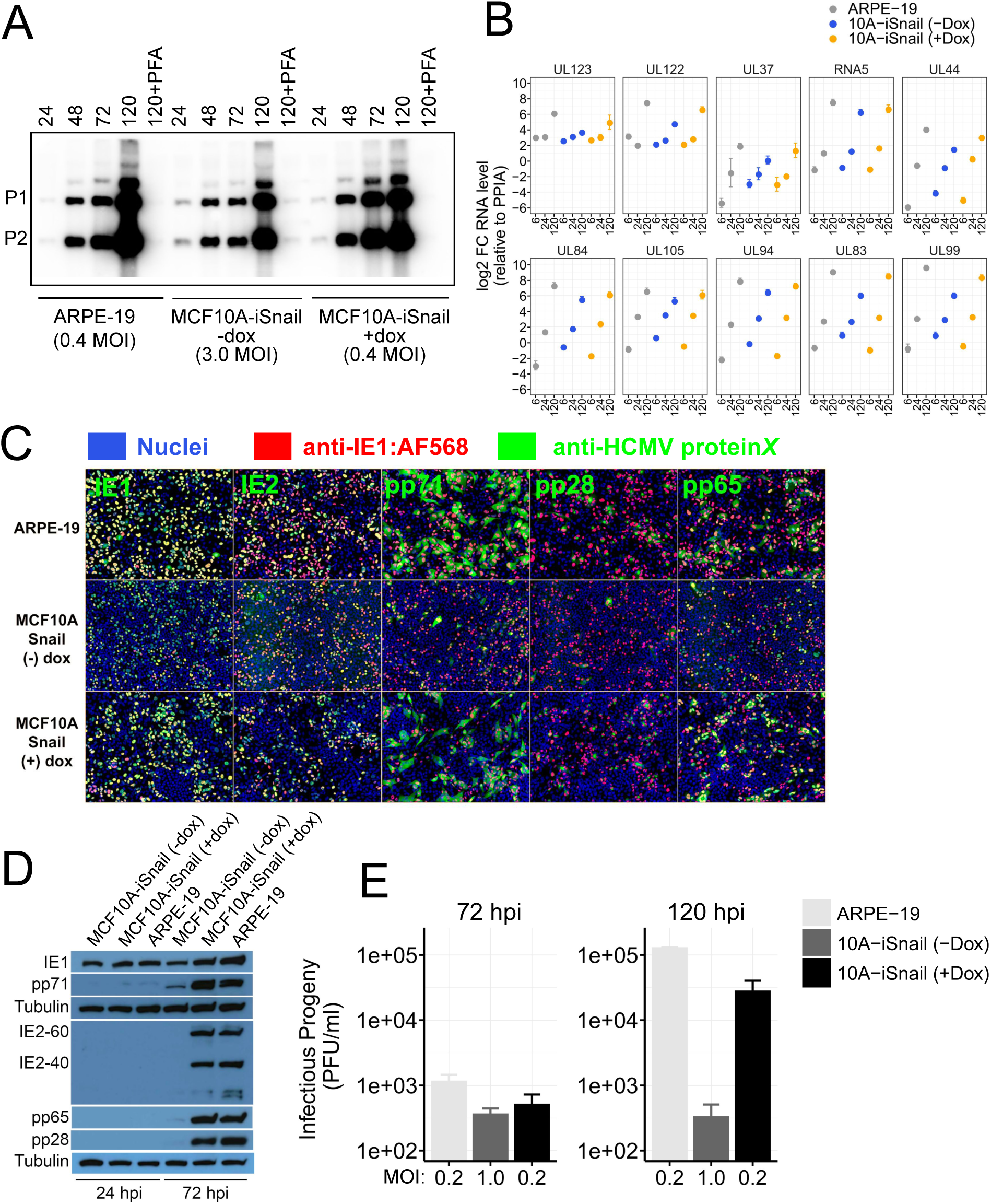
Immunofluorescence detection of HCMV antigens in MCF10A cells with normalized infection conditions at 120 hpi (related to Fig. 6). (A) Southern blot analysis of HCMV viral DNA accumulation after infection with TB40/E-BAC4 using conditions that equalize the number of IE1-expressing cells in each population. ARPE-19, MCF10A-iSnail -dox, and MCF10A-iSnail +dox cells were infected at 0.4, 3.0, and 0.4 MOI, respectively, and total cellular DNA was isolated at the indicated time points for Southern analysis. Cell populations treated with the HCMV polymerase inhibitor Phosphonoformic Acid (PFA) were used as negative controls for DNA replication. (B) Absolute mRNA levels of select viral RNAs across ARPE-19, MCF10A-iSnail -dox, and MCF10A-iSnail +dox cells, at different times post infection, assessed by quantitative RT-PCR. Data points represent the mean fold-change (+/- SD) of each viral transcript compared to the cellular gene PPIA (2^-dCt), from three independent biological replicates assayed in technical triplicate. (C) Immunofluorescence analysis of HCMV proteins in the presence and absence of EMT. Same experimental setup as in Figures 3C and S5, except that the number of IE-1 expressing cells were normalized by using 0.4, 3.0, and 0.4 MOI for ARPE-19, MCF10A-iSnail (-dox), and MCF10A-iSnail (+dox) cells, respectively. Samples were collected at 72 hpi. Similar results were observed at 120 hpi (Figure S8). (D) Western analysis of HCMV proteins at 24 and 72 hpi using normalized infection conditions (0.4, 3.0, and 0.4 MOI for ARPE-19, MCF10A-iSnail -dox, and MCF10A-iSnail +dox cells, respectively). (E) HCMV infectious progeny production at 72 and 120 hpi using infection conditions resulting in approximately 20 % infected cells in each cell type (0.2, 1.0, and 0.2 MOI for ARPE-19, MCF10A-iSnail -dox, and MCF10A-iSnail +dox cells, respectively). Data represent the mean +/- SD of three independent biological replicates per condition.

## Discussion

Herpesviruses cycle through acute and latent infection stages at the organismal level. While immunological control of systemic and local viral spread strongly influences these cycles, latency is believed to also involve the establishment of reversibly permissive states in individual cells. Infection decisions at the single cell level are a consequence of individual cells imprinting their intrinsic biochemical state on the viral genome after entry. Productive cell types provide a biochemical environment supporting all of the biochemical functions required for macromolecular biosynthesis, assembly, and release of new infectious virions. Reversibly permissive cell types support viral entry, but lack a supportive environment for the completion of biosynthesis and assembly; however, dynamic changes to the cellular biochemical environment conditionally enables virion production. For HCMV, reversibly permissivity has been best defined in myeloid progenitor cells, where the intrinsic epigenetic state of undifferentiated progenitor cells imprints on the viral genome, resulting in heterochromatinization of the major immediate-early promoter and restriction of the viral infectious cycle. Cellular differentiation along the macrophage/dendritic-cell lineage, through poorly defined mechanisms, alters the cellular epigenetic state, collaterally de-repressing viral biosynthesis, stimulating virion manufacturing.

Reversible epigenetic silencing of HCMV in myeloid progenitor cells is thought to drive the majority of systemic reactivation of HCMV during periods of immunosuppression. However, it is possible that HCMV resides in latent or alternative states of persistence at additional physiological sites in the body. Epidemiological and histological evidence suggests that epithelial cells in the mammary gland, prostate gland, kidney, and oral cavity could harbor persistent HCMV.

We have found that HCMV can establish a biosynthetically abortive state in mammary epithelial cells that can be conditionally reversed by epithelial-mesenchymal trans-differentiation driven by TGF-β or EMT-stimulating transcription factors. The critical control point limiting HCMV biosynthesis appears to be viral mRNA translation rather than MIEP silencing. Interestingly, a recent study has shown that some HCMV mRNAs utilize the eIF3d-dependent, alternative translation initiation pathway for efficient translation during the late phase of infection^84^. It is possible that EMT potentiates alternative translation initiation pathways, potentiating translation of HCMV mRNAs at late times during infection. Supporting this hypothesis, eIF3D and the alternative scaffolding protein eIF4G2/DAP5 have been shown to be important for translating EMT-related mRNAs (e.g. *SNAIL*, *ZEB1*, and *VIM*) during cancer cell metastasis ^85^. It will be important in future studies to assess the role of alternative translation initiation pathways during HCMV infection in MCF10A epithelial cells.

Several cellular systems where HCMV can enter non-canonical, reversibly permissive biosynthetic states have been described. For instance, experimentally infected NTERA-2 (NT2) germ cell carcinoma cells ^86,87^ or T98G glioblastoma cells ^88–90^ can enter biosynthetically abortive states that can be shifted to permissivity using differentiation factors like retinoic acid or a combination of dibutyryl cAMP and 3-isobutyl-1-methylxanthine. In T98G cells, a latent-like state is thought to be established by heterochromatization of the HCMV MIEP mediated via the cellular histone deacetylase Daxx ^91,92^. The nature of restriction in NT2 cells remains unclear. Our characterization of the effects of EMT-driven trans-differentiation on HCMV biosynthesis in mammary epithelial cells provides an example of a conditionally permissive cell type that interrupts the HCMV infectious cycle at a point downstream of the MIEP. To the best of our knowledge, our results in MCF10A cells provide the first evidence that HCMV biosynthesis can be reversibly modulated in non-transformed epithelial cells by the EMT, a ubiquitous trans-differentiation pathway present in most, if not all, epithelial cells.

We find that HCMV infection and biosynthesis are strongly correlated with mesenchymal cell-state features and that initiating EMT in mammary epithelial cells reverts two nested restrictions limiting HCMV infection and biosynthesis: one involving HCMV entry and one involving viral mRNA translation. Both restrictions can be overcome by initiating EMT before infection, defining the EMT pathway as an important cellular pathway controlling HCMV entry and biosynthesis. Our results support the existence of an unidentified, EMT-sensitive entry mechanism for HCMV into epithelial cells and highlight viral mRNA translation as a potential endogenous cellular restriction point controlling the late phase of HCMV biosynthesis. The EMT pathway could influence HCMV infection *in-vivo* by dynamically enhancing viral entry into epithelial cells and/or, releasing HCMV from a biosynthetically abortive state through translational regulation.

A limitation of our work is that we have only assessed the effects of epithelial-mesenchymal trans-differentiation pathways on HCMV in a small number of epithelial cell lines. Nonetheless, our work in MCF10A mammary epithelial cells establishes that HCMV biosynthesis can be conditionally regulated by stimulating the cellular EMT pathway in epithelial cell lines. Whether this occurs in vivo requires further investigation, presumably in a small animal model of cytomegalovirus infection. However, our results raise the possibility that the epithelial-mesenchymal cell state axis may influence the replication behavior of HCMV under particular circumstances; perhaps contributing to persistence in epithelial tissues, and/or slowly developing pathological conditions linked to HCMV infection, such as cancer.

## Acknowledgments

We thank Zarema Arbieva and Cece Countryman for technical support with RNA-sequencing experiments. We also thank Dr. Alan McLachlan (UIC) for advice, reagents, and technical assistance performing Southern analysis.

---

**Figure S1.**
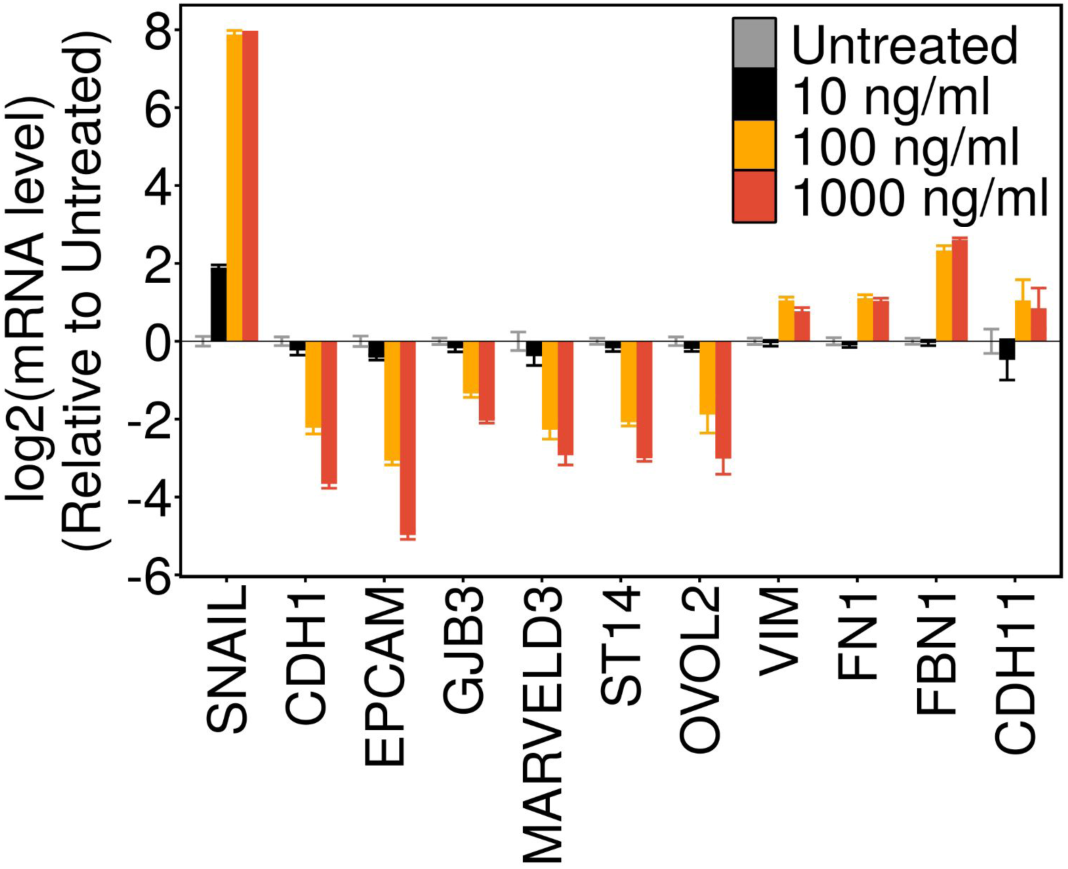
Dose-dependent Induction of an EMT Gene Expression Profile in MCF10A-iSnail Cells (related to Fig. 2). Total RNA was isolated from MCF10A-iSnail cells induced with doxycycline for 2 days and subject to *q*RT-PCR analysis. Increasing concentrations of dox suppress expression of epithelial marker genes and induce expression of mesenchymal marker genes.

**Figure S2.**
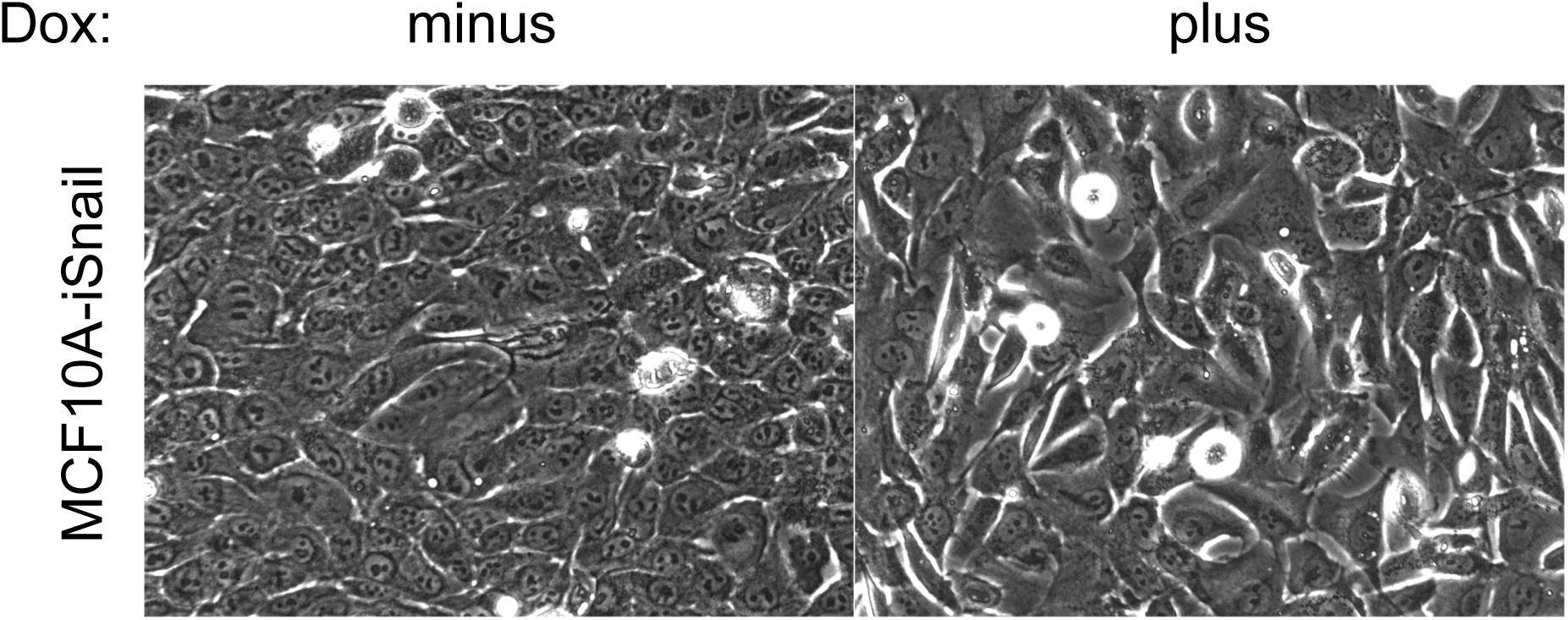
MCF10A-iSnail cells plus and minus doxycycline at high confluency (related to Fig. 2). Typical morphology of induced (+ dox) and uninduced (- dox) cells prior to infection with HCMV, as assessed by phase-contrast microscopy.

**Figure S3.**
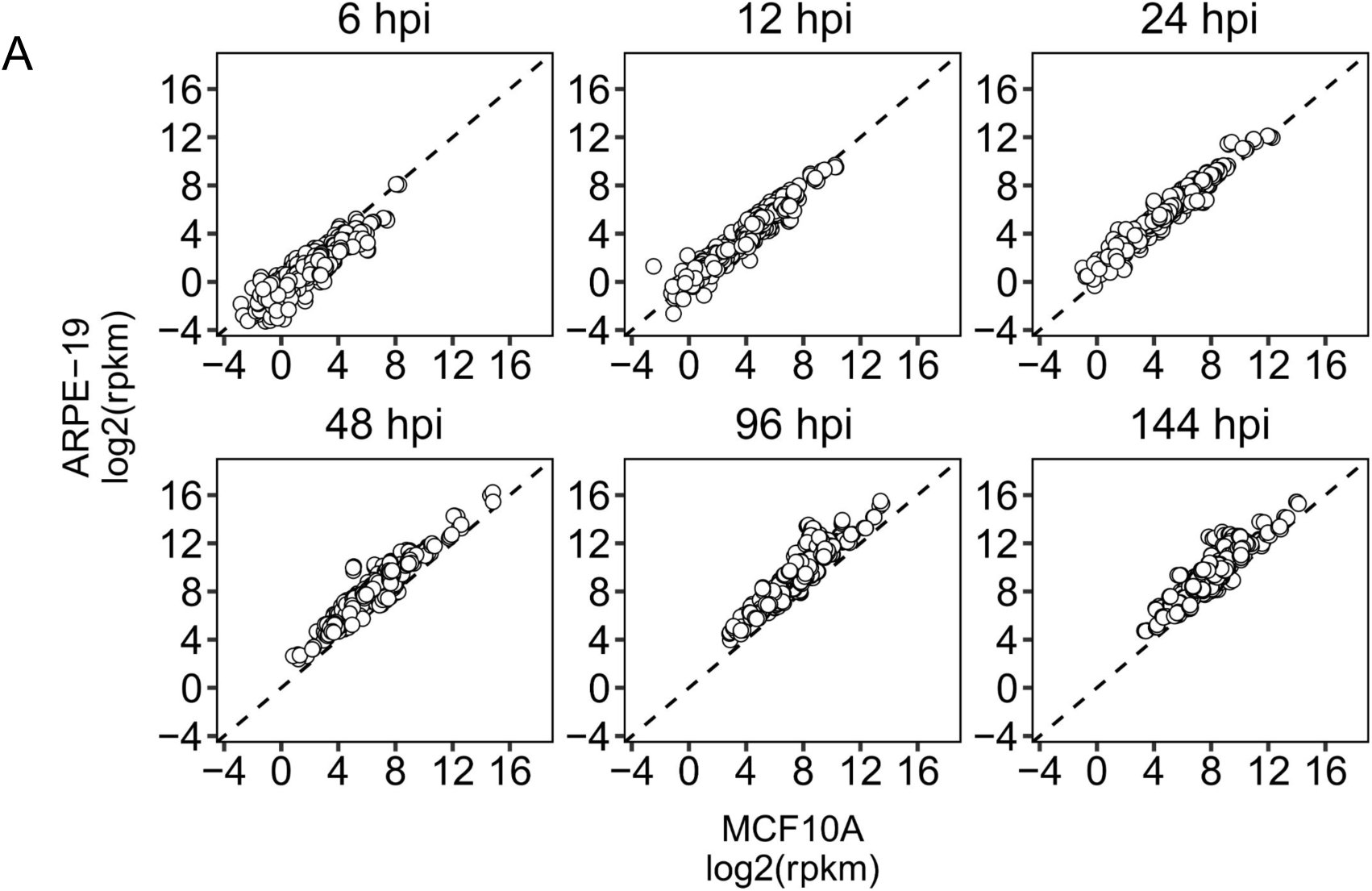
Comparison of absolute expression levels of canonical HCMV RNAs in infected ARPE-19 and MCF10A cells (related to Fig. 3). RNA was extracted from ARPE-19 and MCF10A cells infected with TB40-epi at 6, 12, 24, 48, 96, and 144 hpi. Triplicate infections were performed for each cell type, at each time point, and absolute expression levels were measured by RNA-sequencing (see Methods). Data are visualized as scatter plots. Each point represents the expression level of each of 161 canonical HCMV transcripts from triplicate infections (*n* = 161 x 3 = 483). Triplicate samples were randomly matched across cell types. Dashed line indicates theoretical ratio of 1 (*y* = 1*x* + 0).

**Figure S4.**
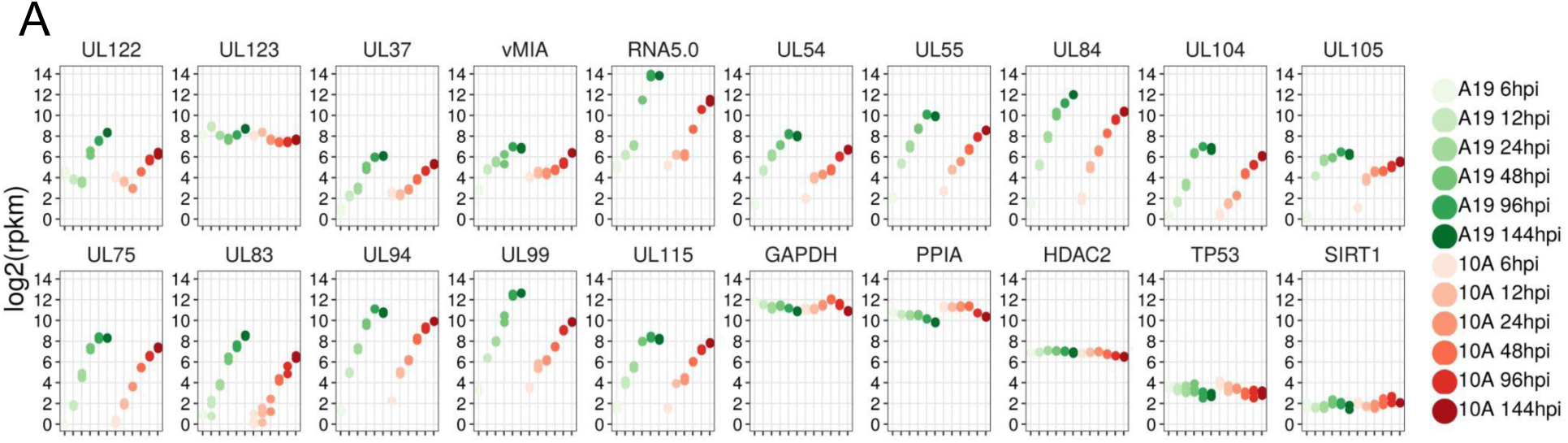
Individual Viral and Cellular Gene Expression Profiles During HCMV Infection in Productive ARPE-19 cells and Restricted MCF10A cells (related to Fig. 3). Mean log2 reads-per-kilobase-per-million of select HCMV and cellular RNAs across time. ARPE-19 and MCF10A cells were infected at a multiplicity of 3 and 5 IU/cell, respectively. Data points represent the geometric mean of three biological replicates per time point. Cellular genes are included to assess cross-sample normalization (GeTMM) and define the dynamic range of RNA expression in cells on the rpkm scale. *SIRT1* and *TP53* represent lowly expressed genes, *HDAC2* a moderately expressed gene, and *PPIA* and *GAPDH* represent highly expressed genes. *hpi*, hours post-infection.

**Figure S5.**
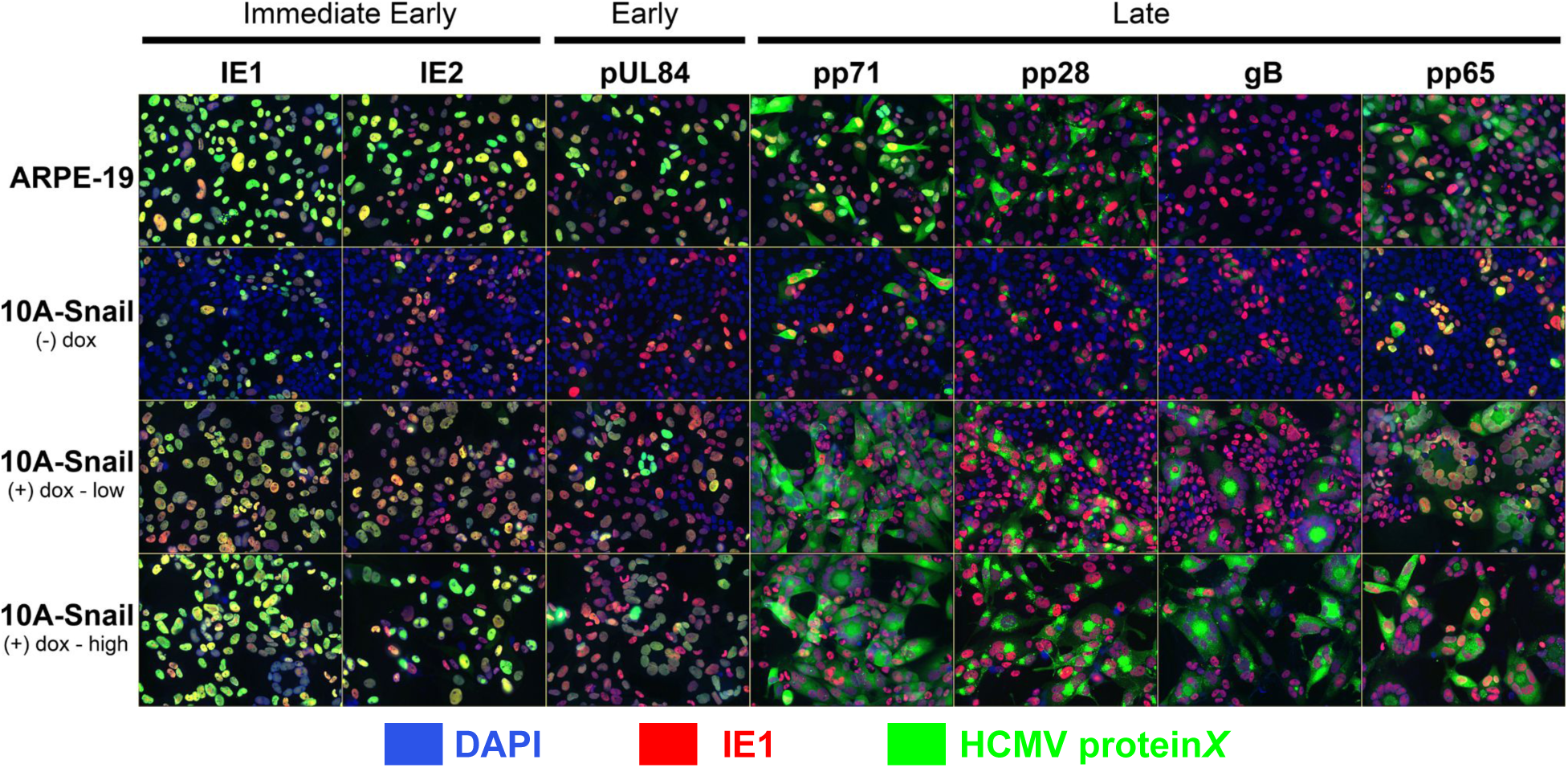
Immunofluorescence detection of HCMV antigens in MCF10A cells, in the presence and absence of EMT, at 120 hpi (related to Fig. 3). MCF10A-iSnail cells were treated with vehicle (- dox), 100 ng/ml doxycycline (+ dox - low), 1 ug/ml doxycycline (+dox - high), infected with HCMV TB40-epi at 1 MOI, and fixed at 120 hpi for immunofluorescence staining with a panel of anti-HCMV antibodies . Cells were counterstained with DAPI (*blue*) to visualize nuclei and an anti-IE1 antibody directly conjugated to CF568 (*red*) to mark infected cells. ARPE-19 cells serve as fully productive control cells. Note the presence of pp65 protein trapped in the nucleus of MCF10A-Snail cells in the absence of doxyclycline/EMT.

**Figure S6.**
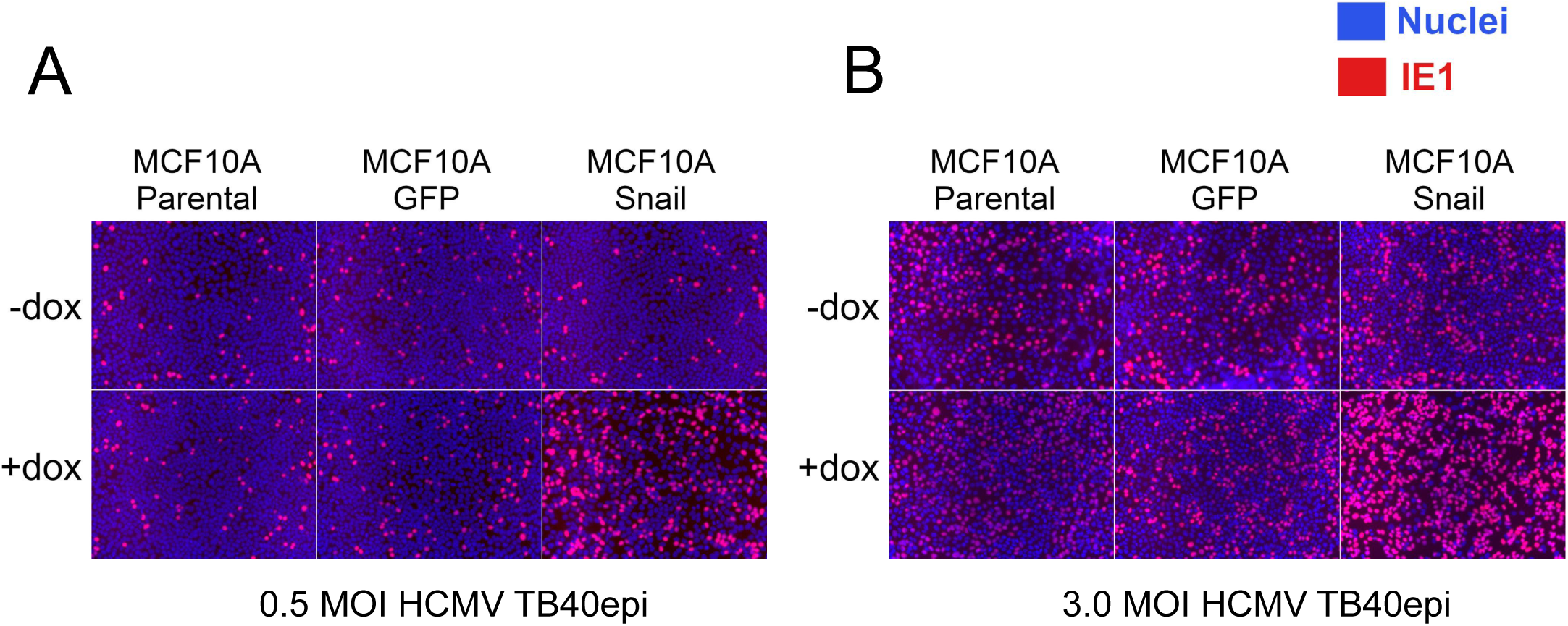
HCMV Entry is Enhanced by EMT (related to Fig. 3). Representative IE1-immunofluorescence images used to quantify HCMV entry in Figures 3D and 3E. Cell populations were infected with HCMV TB40/E-BAC4 prepared in AREP-19 cells at the indicated MOIs and stained for IE1-expression at 16 hpi. The percentage of infected cells was quantified by digitally counting the total number of infected cells in each field (red signal) and dividing by the total number of cells in each field (DAPI/blue signal).

**Figure S7.**
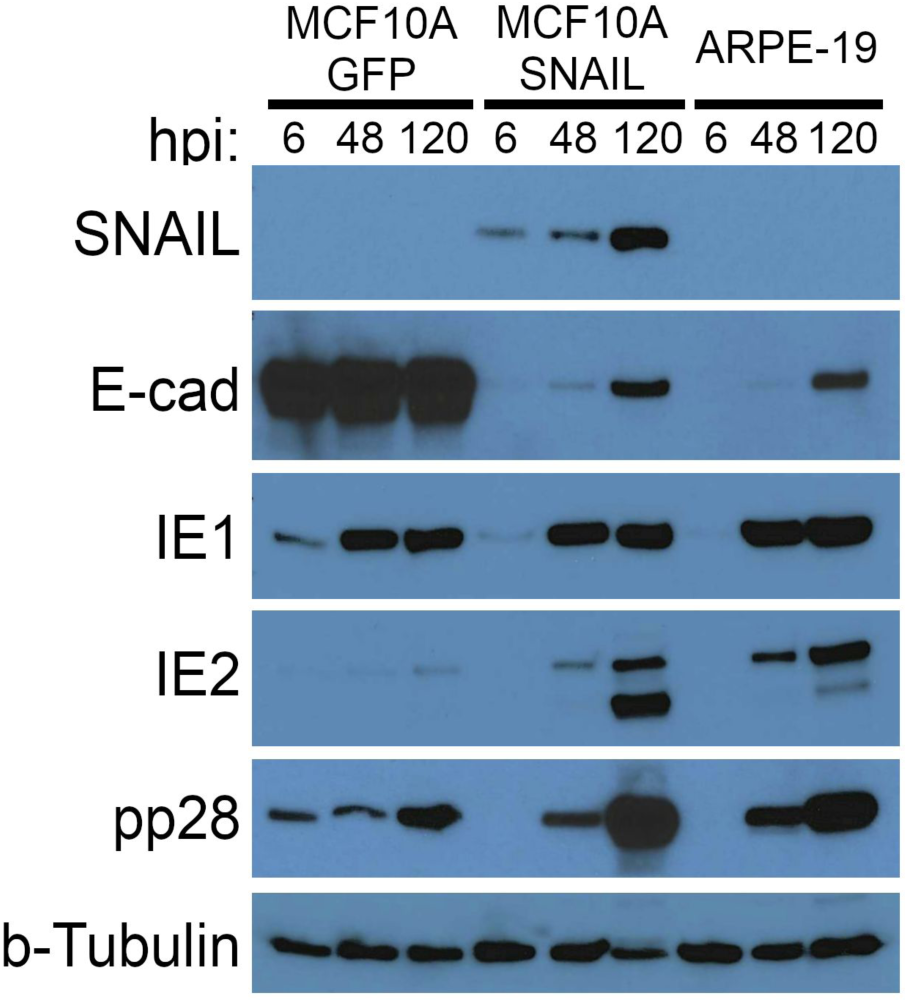
Immunoblot of Cellular and Viral Proteins in MCF10A Cells Constituitively Expressing Snail or GFP and Infected with HCMV (related to Fig. 3). MCF10A cells transduced with lentiviruses expressing Snail or GFP from a constitutive promoter (EF1a) were infected with HCMV TB40/E-BAC4 at 3 or 0.25 IU/cell, respectively, to normalize the percentage of infected cells based on IE1-expression. ARPE-19 cells infected at 0.25 IU/cell were included as productive controls. Snail expression decreases E-cadherin expression and enhances steady-state expression of late IE2 isoforms (i.e. IE2-55 and IE2-40) and pp28; despite IE1 expression being relatively uniform across the cell types. Infection induces steady-state levels of E-cadherin in the mesenchymal cell populations through an unknown mechanism, as previously observed (Oberstein and Shenk, PNAS, 2017).

**Figure S8.**
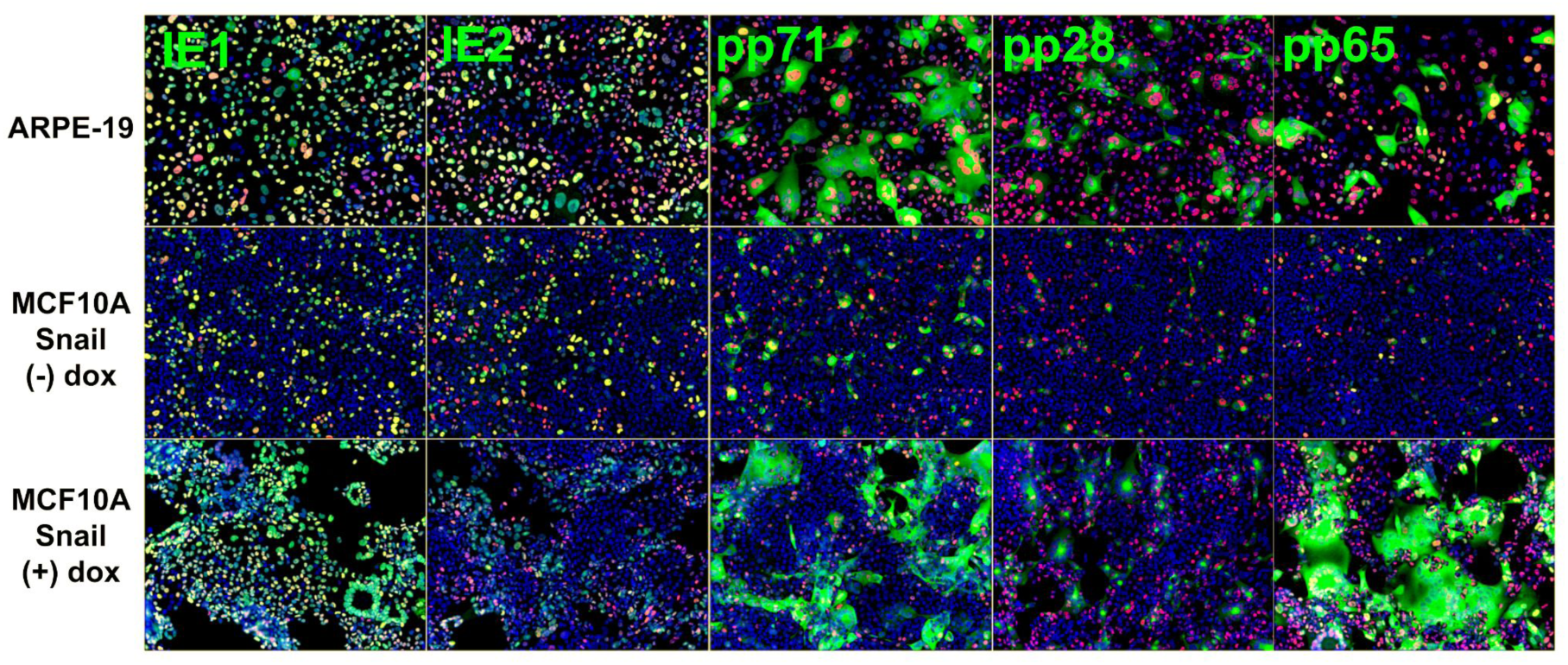
Immunofluorescence detection of HCMV antigens in MCF10A cells with normalized infection conditions at 120 hpi (related to Fig. 5). ARPE-19 or MCF10A-iSnail cells were treated with vehicle (- dox) or 1 ug/ml doxycycline (+dox - high), plated at equal densities, infected with HCMV TB40-epi, and fixed at 120 hpi for immunofluorescence staining with a panel of anti-HCMV antibodies. As in figure 5A, infections were normalized across cell types to equalize the percentage of IE1-positive cells at 24 hpi: 0.4 MOI for ARPE-19 and MCF10A-iSnail (+Dox), 3.0 MOI for MCF10A-iSnail (-Dox). Cells were counterstained with DAPI (*blue*) to visualize nuclei and an anti-IE1 antibody directly conjugated to CF568 (*red*) to mark infected cells.

